# Evidence for a growing population of eastern migratory monarch butterflies is currently insufficient

**DOI:** 10.1101/2020.01.24.918649

**Authors:** Wayne E. Thogmartin, Jennifer A. Szymanski, Emily L. Weiser

## Abstract

The eastern migratory population of monarch butterflies has experienced a multi-decadal decline, but a recent increase in abundance (to 6.05 ha in winter 2018) has led some observers to question whether the population has reversed its long-standing decline and embarked on a trajectory of increasing abundance. We examined this possibility through changepoint analyses, first assessing whether a change in trajectory existed and whether that change was sufficient to alter our estimated risk for the population. We found evidence of a change in trajectory in 2014, but insufficient statistical support for a significantly increasing population since that time (β = 0.285, 95% CI = −0.127, 0.697). If the population estimate for winter 2019 is ≥4.0 ha, we will then be able to credibly assert the population has been increasing since 2014. However, given estimated levels of time series variability, presumed habitat capacity and no recent change in status or trend, there was a 13.5% probability of observing a population estimate as large or larger than was reported for winter 2018. Despite insufficient evidence for an increasing population, near-term risk of quasi-extinction by 2023 has declined (mean risk declining from 43% to 20%) because of higher abundance estimates since 2014. Our analyses highlight the incredible difficulty in drawing robust conclusions from annual changes in abundance over a short time series, especially for an insect that commonly exhibits considerable year-to-year variation. Thus, we urge caution when drawing conclusions regarding species status and trends for any species for which limited data are available.

---

> “Short-term fluctuations may or may not contain messages about longer-term trends” – Art Shapiro

Populations vary over time in their abundance, and this variability can impart uncertainty to the status and trend of a species. As population dynamics approach extinction, dynamics become more variable (Holmes and Fagan 2006), which means short-term highs might become higher, even while abundance is declining on average. In addition to the stochastic variation in abundance imposed by the environment, uncertainty in species status and trend arises from population sizes most often being estimated rather than counted; trends being inferred from limited duration time series; and latent characteristics of a population, such as its relation to carrying capacity or quasi-extinction thresholds, generally being inferred properties rather than an observable quantity. Thus, given these various sources of uncertainty, it is difficult enough to determine the trajectory for a population, let alone any change that may occur in that trajectory, especially one that may occur near the terminus of a time series based on limited data.

Estimates of the population size of the eastern North American migratory population of monarch butterflies (*Danaus plexippus*, hereafter monarchs) in their overwintering locations in high-elevation oyamel fir (*Abies religiosa*) forests of central Mexico suggest a long-term decline in abundance. Using a model allowing separation of observation-induced error from natural process variability, Semmens et al. (2016) estimated monarchs declined by 84% between the winters beginning in 1996 (18.19 ha) and 2014 (0.67 ha), with an estimated annual population rate of change of 0.94. This rapid decline in monarch abundance led to widespread concern regarding the imperilment of the species (Brower et al. 2011), including a petitioning of the U.S. Fish and Wildlife Service (USFWS) to consider listing the species under the U.S. Endangered Species Act (ESA) of 1973 (Center for Biological Diversity et al. 2014).

The estimated rate of decline (λ = 0.94) in monarchs was, however, considerably uncertain, with credible intervals spanning from as low as 0.69 to as high as 1.30. This uncertainty, in turn, led to considerable uncertainty in the estimates of risk faced by the population; for instance, depending on the quasi-extinction threshold chosen, the range of uncertainty in the risk was as much as one or two orders of magnitude wide (i.e., 0-34% at a 0.01 ha quasi-extinction threshold and 7-88% at 0.25 ha). The principal reasons for this large uncertainty in the trajectory of monarchs and their subsequent risk of further decline are the environmental and biological variability this insect faces over its annual cycle and our ability to intuit the species response to this variability with the limited data available from monitoring programs. Density-independent mortality, caused by a wide array of annually variable environmental stressors, is offset against density-dependent reproduction (Yakuba et al. 2004, Drury and Dwyer 2006, Flockhart et al. 2012, Marini and Zalucki 2017), and this tension between birth and death processes plays out over multiple generations and across the vastness of eastern North America (Flockhart et al. 2015, Oberhauser et al.2017). In some years, these processes complement one another, leading to booms or busts in the population (Himes Boor et al. 2018). In other years, increases in one are offset by the other, mitigating any sizeable year-to-year change in population size.

In winter 2018, estimates of monarch abundance in their overwintering areas indicated monarchs increased by 144% over their previous year’s abundance, to an index of population size of 6.05 ha (Conanp and WWF-Mexico 2019). This estimate has led some observers to question whether the population has grown in recent years to the point at which it is no longer at risk. This seemingly simple question is manifold in nature. The question suggests that there may have been a change in the trajectory of the species in recent years, from a population in decline to one of increase, that in turn begs whether the evidence of this change in trajectory supports a reduced risk of quasi-extinction. An alternative possibility could be that the underlying status and trajectory of the population had not changed but instead the species demonstrated the extreme variability in year-to-year abundance that is not uncommon for insects.

To address this question, we conducted a time-series analysis examining whether the observed series of population sizes experienced changes in mean or trajectory anywhere over the 25-year period of record. We also refit the model of Semmens et al. (2016) with subsequent years of data reflecting the most recent observations of population size. In these ways, we can test whether the population has reversed its long-term decline and the risk it faces has appreciably declined as a result. The population as measured in Mexico reached its nadir in abundance in winter 2013 (WWF 2016); we hypothesized that any change in status and any reversal of trend should occur at this point in the time series.

## Methods

The overwinter index of population size (in hectares) we used in our models was that used by the USFWS in its Species Status Assessment for informing considerations of whether listing under the ESA is warranted. With these data, we evaluated two models, a step arrangement 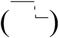 evaluating whether there was a demonstrable change in status (i.e., mean abundance) during the time period and a segmented model 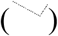 examining whether there was a change in the trend; we specifically tested for a reversal of trend from a period of decline to one of growth. We fit these models in R (R Core Team 2018) with the *changepoint* (Killick et al. 2016) and *chngpt* (Fong and Gilbert 2017) packages. Assumptions of independent, normally distributed data (on a log_e_ scale) with constant variance pre- and post-change were evaluated with Shapiro and Kolmogorov-Smirnov tests and inspection of quantile-quantile and autocorrelation plots.

Pleasants (2017) suggested there was sufficient milkweed in the upper midwestern U.S. to support a mean population size overwintering in Mexico of 3.2 ha. He also asserted that in some years, the reported abundance is likely to be lower because of the accumulation of poor conditions faced by the population during its annual cycle, whereas in some years favorable conditions will lead to a population increase higher than 3.2 ha. We calculated the probability from a log-normal distribution of observing a 6.05-ha population relative to the 3.2-ha expected population size. We calculated the variance for this log-normal distribution from the variance of the post-2013 period.

Given that a changepoint was identified and the post-changepoint period was nonsignificantly increasing (95% confidence interval of the slope parameter overlapping 0) (see Results), we asked the question: How many more years of positive increase would be necessary to provide statistically robust evidence that the population was growing? To evaluate this question, we extrapolated the post-changepoint period abundance given the estimated post-changepoint slope and refit the segmented changepoint model with additional years of extrapolated abundance.

To examine potential change in risk of quasi-extinction given the recent population size estimates for monarchs, we re-fit the Bayesian state-space model of Semmens et al. (2016), calculating a new measure of risk with the additional years of estimated abundance since winter beginning in 2014 (the last winter included in Semmens et al. 2016) (Figure 1). We set the quasi-extinction threshold to 0.25 ha. We calculated this risk to winter 2023 (+5 years) and compared it to the risk estimate for 2023 from the original Semmens et al. (2016) calculation. An important aspect of the Semmens et al. model is that it used the conjoint patterns in overwinter abundance and egg production in the breeding grounds to disentangle observation error from process error. This egg production information is derived from an extrapolation of egg density data applied to year-specific estimates of monarch-appropriate land cover. Unfortunately, annual land cover information is insufficiently compiled for the most recent years at this time to reconstitute a longer time series of egg production; nevertheless, the egg density time series used in Semmens et al. is sufficient for the purposes of separating the two principal sources of error. An important assumption we made in re-estimating risk was that there was no change in the trajectory of the population over the full time series.

**Figure 1.**
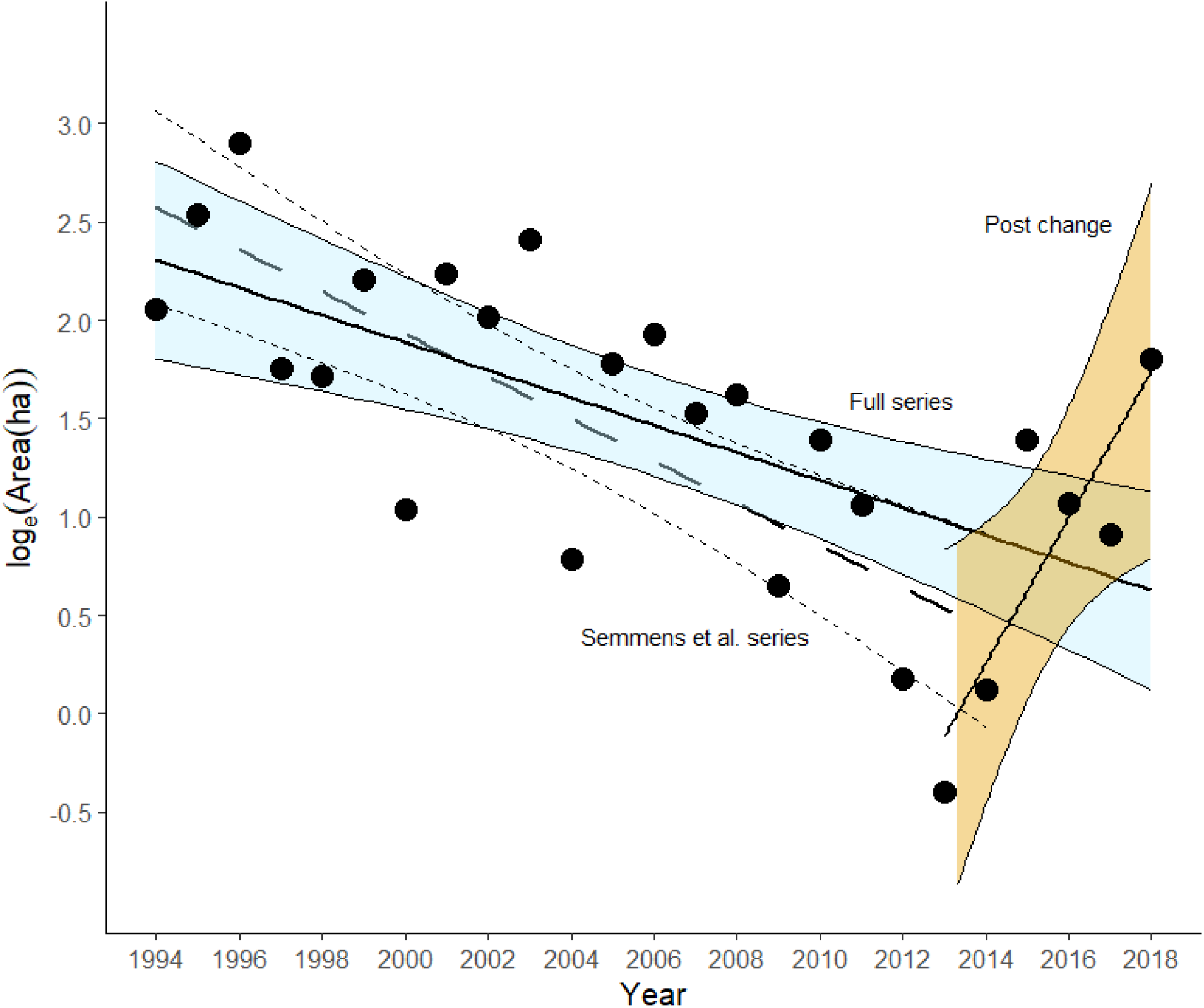
Time series segmented into original series analyzed by Semmens et al. (2016)(1994-2014), the full set (1994-2018), and the post-change period (2013-2018). Risk of quasi-extinction was calculated for each of these three periods.

The time series after the estimated change point (described below) is too short to allow proper estimation using the methods of Semmens et al. for only that post-changepoint period. Thus, we used a simpler population-projection population viability analysis (Morris and Doak 2002) to estimate risk for the post-changepoint period in year 2023. As before, we set the quasi-extinction threshold to 0.25 ha, counting the number of 1,000 simulations of population size dropping below that threshold, and then compared that proportion to the quasi-extinction risk estimated by Semmens et al. (2016). Calculation of this population-projection population viability analysis, with 95% bootstrapped confidence intervals, was conducted following Morris and Doak (2002, box 7.3) with the *popbio* package using the countCDFxt function (Stubben et al. 2018) in R.

## Results

When examining the time series of overwinter abundance of the eastern migratory population of monarch butterflies for a change in mean abundance (i.e., step change), we identified a single credible changepoint in winter 2009. For the period preceding this year, mean abundance was 6.69 ha (95% CI = 4.43, 8.94). For the period after winter 2009, mean abundance was 1.52 ha (95% CI = >0.001, 4.68). The population variance was 15% higher in this latter period (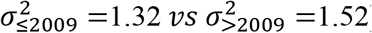), exhibiting greater variability at lower population sizes. If the underlying milkweed is currently sufficient to support a winter population of 3.2 ha (Pleasants 2017), then a population as large or larger than 6.05 ha is expected to occur 13.5% of the time.

Fitting a segmented changepoint model, rather than a step function, suggested the best-supported year for the changepoint threshold was 2014 (likelihood ratio λ = 8.167, *p* = 0.0221; bootstrapped 95% CI = 2002, 2026), with 2013 close behind (Figure 2). The slope describing the decline of monarchs in the period before winter 2014 was −0.103 (Table 1), whereas after this winter the population exhibited a non-significant increase, though with confidence intervals >5:1 in favor of an increase (β = 0.285, 95% CI = −0.127, 0.697) (Figure 1).

**Table 1.**
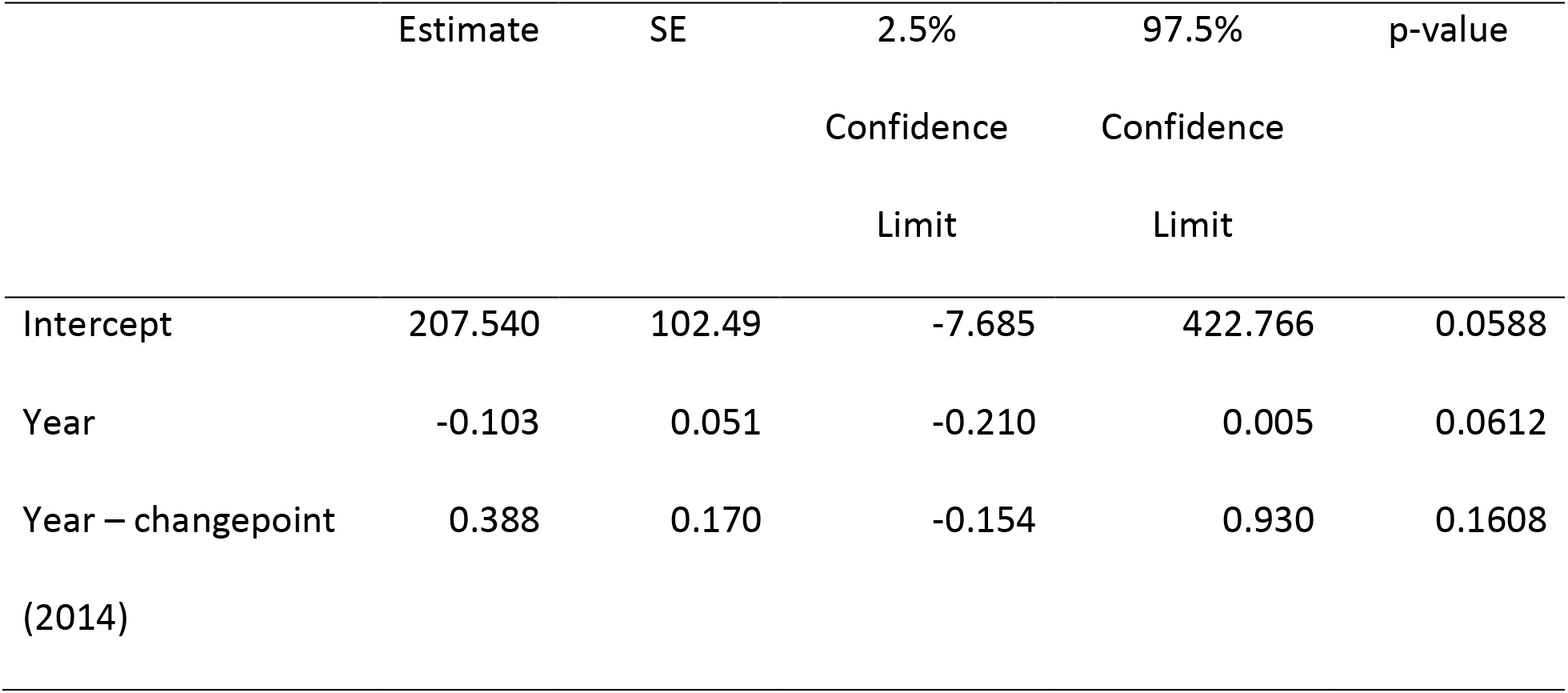
Parameter estimates for the best-supported linear segmented changepoint model for 1994-2018 estimates of overwinter abundance of the eastern migratory monarch butterfly population.

**Figure 2.**
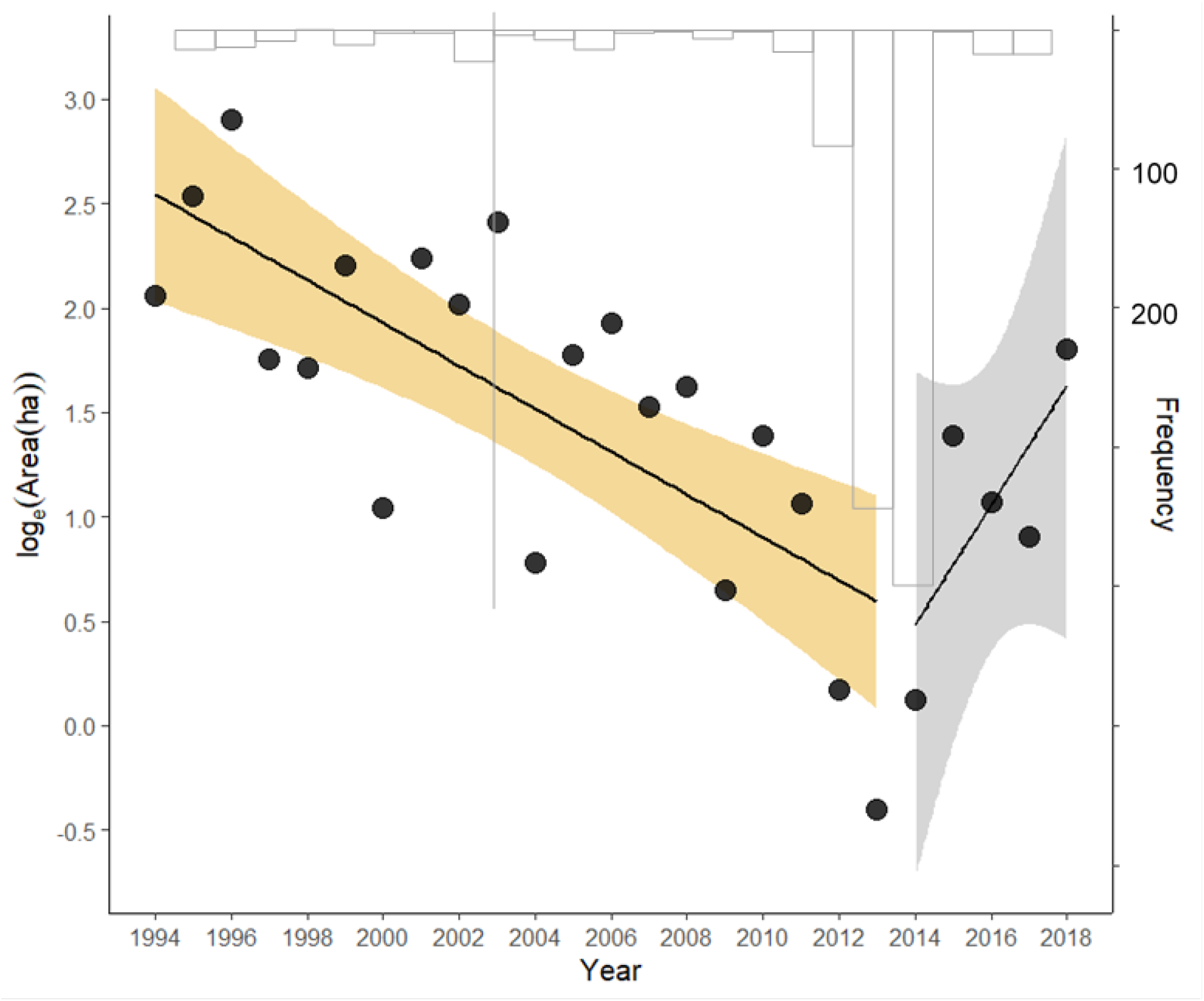
Segmented time series of the index of overwinter abundance (ha) of eastern migratory monarch butterflies. The bootstrapped frequency of the changepoint estimate from 10^3^ replicates is provided (inverted, in gray); the gray line represents the lower 2.5% symmetric bootstrap confidence limit.

Residuals from these step and segmented models before and after their changepoints were independent, normally distributed about their respective mean, and had constant variance. Comparing the segmented model (AIC = 45.3) with the step model (AIC = 49.5) suggested an 88% probability (odds 7.2:1) that the segmented model served as a better description of the data. Both models were appreciably better than a null model comprising only an intercept (AIC = 62.8) and a linear model regressing the log_e_(overwinter estimate) against year (AIC = 51.5).

If the winter 2019 population continues the mean rate of increase observed since 2014, then with this single additional year of data, we would have sufficient information statistically to conclude the population was growing (*β* = 0.399, 95% CI = 0.072, 0.727). Further, if the index of abundance was any value ≥4.00 ha, this amount too would be statistically sufficient (*p* < 0.05) to support a conclusion that the population was growing. Any value <4.00 ha, however, would cast doubt on a growing population.

Refitting the model of Semmens et al. (2016) to assess whether the population’s risk of quasi-extinction had appreciably changed, as measured over the entire 25-year period of record, indicated declining risk with improving abundance assuming no change in trajectory (Table 2). If, however, the post-2014 winter trajectory continues, the risk of quasi-extinction by 2023 could drop to negligible levels.

**Table 2.**
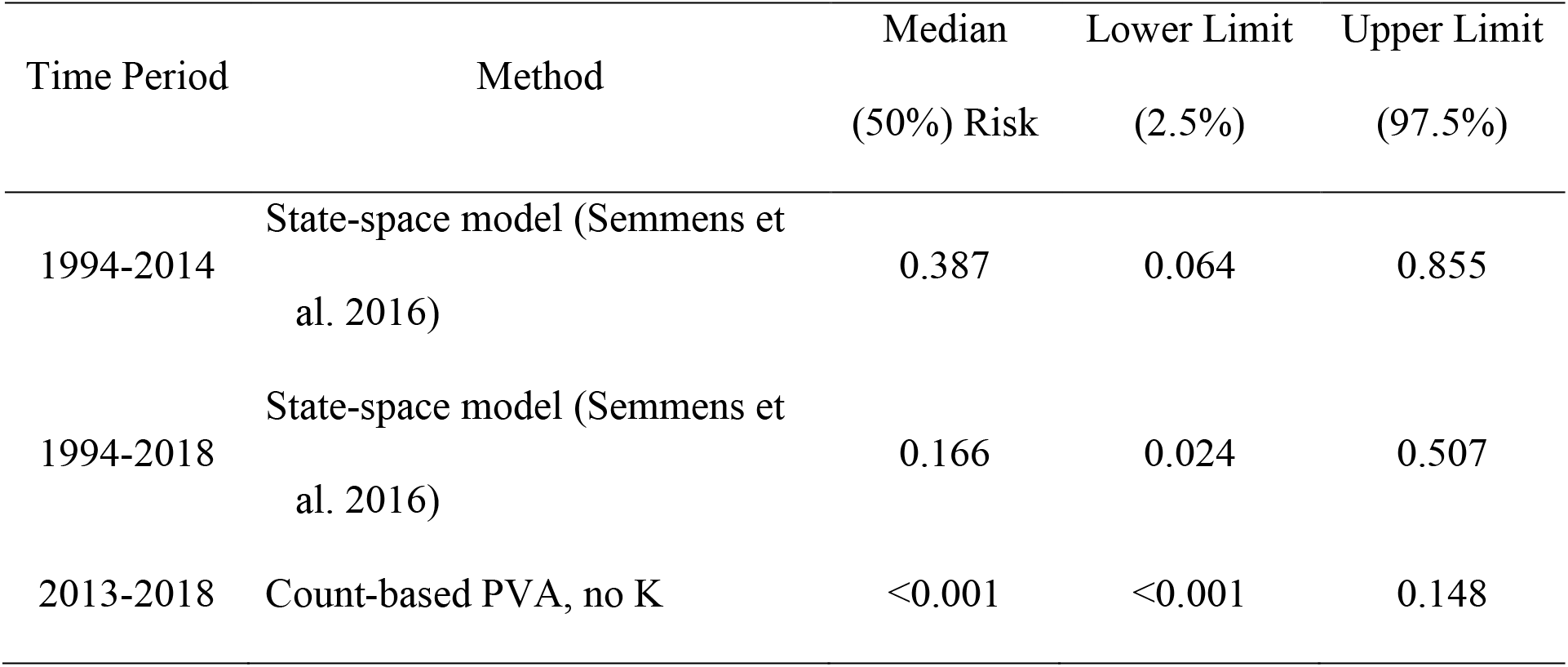
Median risk of quasi-extinction by 2023 (quasi-extinction threshold = 0.25 ha) from the original Semmens et al. (2016) model (with overwinter abundances from 1994-2014), for the full time series of overwinter abundances (1994-2018, again using the methods of Semmens et al.), and for the post-change period (2013-2018), using a count-based population viability model.

## Discussion

At this time, there is insufficient statistical evidence to confidently assert that the eastern migratory monarch population has significantly grown since winter 2014. If the dynamic of population growth for the few years post-winter 2014 holds, then winter 2019-2020’s population size estimate should provide evidence as to whether the trend has credibly changed from one of decline to one of increase.

In a noisy time series, stochastic fluctuations may lead to observed increases over relatively long periods, even when populations have an average negative growth rate. Similarly, stochastic fluctuations may cause a population to decrease, even when the long-term average growth rate is positive. Our analysis and the uncertainty it reveals highlights the difficulty in assessing species status and trend with even a 25-year dataset, especially when interannual variation is high. Semmens et al. (2016) reported a mean declining dynamic, but one with a non-negligible probability of a possible underlying growth rate that was positive. Their findings showed that two-thirds of the credible interval distribution about their estimate of the population growth rate was <1, indicating that the odds were 2:1 in favor of a declining population. Nevertheless, one-third of the distribution suggested a stable or growing population. Conversely, based on the interval width we calculated for the post-2014 trajectory, the odds are roughly 5:1 in favor of an increasing population. Unfortunately, the post-2014 period is too short to confidently conclude, at this time, a reversal in trajectory.

Despite a lack of statistical support for a positive trend since 2014, the near-term risk of quasi-extinction appears to have declined by at least half. Nevertheless, there remains a 1 in 5 chance of quasi-extinction within 5 years if the population has not truly changed trajectory. If it has changed trajectory, then the near-term risk is further ameliorated to near zero by 2023 (also see table 2, Semmens et al. 2016).

In any time series, the sample size is the number of years, and 10-30 years are often necessary to detect a significant trend even for species with average interannual variation (Urquhart, 2012; White, 2018). Despite the challenge of high interannual variation, the monarch butterfly is an iconic and highly visible species that benefits from strong public interest (Diffendorfer et al., 2014) and a corresponding availability of data (Ries and Oberhauser, 2015). For many species considered for listing under the ESA, even less information is available for evaluating the statistical support for any apparent decline. Thus, the challenge of assessing trend becomes even greater as one examines short-term time series or smaller periods of time within long-running time series; what may initially appear to be a short-term trend may have no statistical support in the context of the population’s history. While assessing subsets of a time series could be a useful way to evaluate whether a species is moving toward recovery, caution is warranted when making conclusions based on limited data. Aside from the estimate of trend, other metrics can be useful in such cases, such as whether mean abundance falls below the estimated threshold for a secure population. In the case of the monarch butterfly, the recent mean of 1.52 ha falls well below the threshold of 6.0 ha estimated by Semmens et al. (2016) and established by the three nations of Canada, U.S. and Mexico as the near-term population goal for the eastern population of migratory monarch butterflies. If we take this 6.0 ha threshold as a recovery criterion and assume a 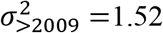, then the population is likely to need to reach a mean of 6.85 ha for 3 years to confidently assert the population has crossed this threshold (analysis not shown). Thus, this mean population size warrants continuing concern given the uncertain growth in recent years and the high year-to-year variability exhibited by this insect species.

## Acknowledgments

Any use of trade, firm, or product names is for descriptive purposes and does not imply endorsement by the US Government. Views expressed in this article do not necessarily represent those of the US Fish and Wildlife Service. We appreciate M. Post Ven Der Burg and two reviewers for comments made on an earlier version of this manuscript.

## Author Contributions

WT and JS conceived the study, WT conducted the analyses, and WT, JS, and EW wrote the manuscript.

