## Supplementary material for "Evidence for a growing population of eastern migratory monarch butterflies is currently insufficient": R Code

if(!require(changepoint)){
  install.packages('changepoint')
}

library(changepoint)
if(!require(changepoint.np)){
  install.packages('changepoint.np')
}

library(changepoint.np)


# y<-log(c(7.81,12.61,18.19,5.77,5.56,8.97,2.83,9.36,7.54,11.12,2.19,5.91,
#          6.87,4.61,5.06,1.92,4.02,2.89,1.19,0.67,1.13,4.01,2.91,2.48,6.05))
# 
# yalt<-log(c(7.81,12.61,18.19,5.77,5.56,9.05,2.83,9.35,7.54,11.12,2.19,5.92,
#              6.87,4.61,5.06,1.92,4.02,2.89,1.19,0.67,1.13,4.01,2.91,2.48,6.05))

###Official estimates, from ES (J. Szymanski)
y<-log(c(7.81,12.61,18.19,5.77,5.56,9.05,2.83,9.36,7.54,11.12,2.19,5.91,
         6.87,4.61,5.06,1.92,4.02,2.89,1.19,0.67,1.13,4.01,2.91,2.48,6.05))
length(y)
MASS::truehist(y, n=5) #roughly normal on the log-scale

year <-seq(1994,2018)

monarchdata <- cbind(year, y)

monarchts <- ts(y, frequency=1, start=1994, end=2018)
#str(monarchts)
#View(monarchts)
plot(monarchts)

Box.test(monarchts, lag=2, type="Ljung-Box")
kpss.test(monarchts, null="Trend")


###Is there a changepoint in the mean
monarchts.amoc2 <-
  cpt.mean(
    monarchts,
    penalty = "SIC",
    pen.value = 0,
    method = "AMOC",
    Q = 5,
    test.stat = "Normal",
    class = TRUE,
    param.estimates = TRUE,
    minseglen = 1
  )
monarchts.amoc3 <-
  cpt.mean(
    monarchts,
    penalty = "BIC",
    pen.value = 0,
    method = "AMOC",
    Q = 5,
    test.stat = "Normal",
    class = TRUE,
    param.estimates = TRUE,
    minseglen = 1
  )
monarchts.amoc3a <-
  cpt.mean(
    monarchts,
    penalty = "BIC",
    pen.value = 0,
    method = "AMOC",
    Q = 5,
    test.stat = "Normal",
    class = FALSE,
    param.estimates = TRUE,
    minseglen = 1
  )
monarchts.amoc3b <-
  cpt.mean(
    monarchts,
    penalty = "BIC",
    pen.value = 0,
    method = "SegNeigh",
    Q = 5,
    test.stat = "Normal",
    class = TRUE,
    param.estimates = TRUE,
    minseglen = 1
  )
monarchts.amoc4 <-
  cpt.mean(
    monarchts,
    penalty = "AIC",
    pen.value = 0,
    method = "AMOC",
    Q = 5,
    test.stat = "Normal",
    class = TRUE,
    param.estimates = TRUE,
    minseglen = 1
  )
monarchts.amoc5 <-
  cpt.mean(
    monarchts,
    penalty = "Hannan-Quinn",
    pen.value = 0,
    method = "AMOC",
    Q = 5,
    test.stat = "Normal",
    class = TRUE,
    param.estimates = TRUE,
    minseglen = 1
  )
monarchts.amoc6 <-
  cpt.mean(
    monarchts,
    penalty = "Asymptotic",
    pen.value = 0.5,
    method = "AMOC",
    Q = 5,
    test.stat = "Normal",
    class = TRUE,
    param.estimates = TRUE,
    minseglen = 1
  )

#Conclusion, by and large, the penalty has no consequence to the result
cpts(monarchts.amoc3) #All are equivalent, suggesting a change in year 15 of the time series
param.est(monarchts.amoc3)
plot(monarchts.amoc3, lwd=3, xlab="Year", ylab="log(Area[ha])")

summary(monarchts.amoc3)
logLik(monarchts.amoc3)


###Is there a changepoint in the variance
monarchtsv1.BIC <- cpt.var(monarchts, method='AMOC',penalty='BIC',pen.value='2*log(n)')
cpts(monarchtsv1.BIC)
param.est(monarchtsv1.BIC)

plot(monarchtsv1.BIC,cpt.width=3)


###Is there a change in the mean AND variance
monarchtsmv1.segneigh <- cpt.meanvar(monarchts,test.stat='Normal',method='SegNeigh',Q=4,penalty="SIC")
cpts(monarchtsmv1.segneigh)
param.est(monarchtsmv1.segneigh)
plot(monarchtsmv1.segneigh,cpt.width=3,cpt.col='blue')

monarchtsmv1.binseg <- cpt.meanvar(monarchts,test.stat='Normal',method='BinSeg',Q=4,penalty="SIC")
cpts(monarchtsmv1.binseg)
param.est(monarchtsmv1.binseg)
plot(monarchtsmv1.binseg, lwd=3, xlab="Year", ylab="log(Area[ha])",cpt.width=3,cpt.col='blue')

exp(param.est(monarchtsmv1.binseg)$mean[1])
exp(param.est(monarchtsmv1.binseg)$mean[2])

exp(param.est(monarchtsmv1.binseg)$variance[1])
exp(param.est(monarchtsmv1.binseg)$variance[2])

norm.interval = function(data, variance = var(data), conf.level = 0.95) {
   z = qnorm((1 - conf.level)/2, lower.tail = FALSE)
   xbar = mean(data)
   sdx = sqrt(variance/length(data))
   c(xbar - z * sdx, xbar + z * sdx)
}

norm.interval(exp(param.est(monarchtsmv1.binseg)$mean[1]), 
              exp(param.est(monarchtsmv1.binseg)$variance[1]))


norm.interval(exp(param.est(monarchtsmv1.binseg)$mean[2]), 
              exp(param.est(monarchtsmv1.binseg)$variance[2]))


###how likely is a 6.05 ha population?
m <- param.est(monarchtsmv1.binseg)$mean[2]   #exp(param.est(monarchtsmv1.binseg)$mean[2])
s <- sqrt(param.est(monarchtsmv1.binseg)$variance[2])
location <- log(m^2 / sqrt(s^2 + m^2))
shape <- sqrt(log(1 + (s^2 / m^2)))
print(paste("location:", location))
print(paste("shape:", shape))
plnorm(log(6.05), location, shape)


draws <- rlnorm(1000, location, shape)
v <- log(6.05)
plot(density(draws), main="")
abline(v=v, lty=2, col="black")

###Relative to John Pleasants' estimate of the population size under mean conditions for the amount of milkweed expected to be in the landscape
m32 <- log(3.2)  #if we assume there is an underlying 3.2 ha mean equivalent milkweed
s <- sqrt(param.est(monarchtsmv1.binseg)$variance[2])
location32 <- log(m32^2 / sqrt(s^2 + m32^2))
shape32 <- sqrt(log(1 + (s^2 / m32^2)))
print(paste("location:", location32))
print(paste("shape:", shape32))
1-plnorm(log(6.05), location32, shape32)

draws32 <- rlnorm(1000, location32, shape32)
v <- log(6.05)
plot(density(draws32), main="")
abline(v=v, lty=2, col="black")


##evaluating assumptions for step changepoint
cpt.binseg <- cbind(c(0,cpts(monarchtsmv1.binseg)), seg.len(monarchtsmv1.binseg))
data.binseg <- data.set(monarchtsmv1.binseg)
data <- data.binseg
shapiro.func=function(x){
  out=shapiro.test(data[(x[1]+1):(x[1]+x[2])])
  return(c(out$statistic,p=out$p.value))}
apply(cpt.binseg,1,shapiro.func)

ks.func=function(x){
  tmp=data[(x[1]+1):(x[1]+x[2])]
  out=ks.test(tmp,pnorm,mean=mean(tmp),sd=sd(tmp))
  return(c(out$statistic,p=out$p.value))}
apply(cpt.binseg,1,ks.func)

qqnorm.func=function(x){
  qqnorm(data[(x[1]+1):(x[1]+x[2])])
  qqline(data[(x[1]+1):(x[1]+x[2])])}
out=apply(cpt.binseg,1,qqnorm.func)

acf.func=function(x){
  acf(data[(x[1]+1):(x[1]+x[2])])}
out=apply(cpt.binseg,1,acf.func)

means=param.est(monarchtsmv1.binseg)$mean
m1.resid=data-rep(means,seg.len(monarchtsmv1.binseg))
shapiro.test(m1.resid)
ks.test(m1.resid,pnorm,mean=mean(m1.resid),sd=sd(m1.resid))

qqnorm(m1.resid)
qqline(m1.resid)
acf(m1.resid)


###fitting the step model with library(chngpt)
library(chngpt)
monarchdata.df <- as.data.frame(monarchdata)
monarch.step <-
  chngptm(
    formula.1 = y ~ 1,
    formula.2 =  ~ year,
    monarchdata.df,
    type = "step",
    family = "gaussian",
    est.method = "grid",
    var.type = "bootstrap",
    save.boot = TRUE
  )
summary(monarch.step)
#str(monarch.step)  #AIC = 49.5
monarch.step$logliks[14]


monarch.step$best.fit$aic

AIC(lm(y~1, monarchdata.df))
AIC(lm(y~year, monarchdata.df))

###continuous threshold linear regression
#library(chngpt)
monarch.fit <- chngptm (formula.1= y~1, formula.2=~year, monarchdata.df,
             type="segmented", family="gaussian", est.method="fastgrid", var.type="bootstrap", save.boot=TRUE)
summary(monarch.fit)
str(monarch.fit)
est <- lincomb(monarch.fit, comb=c(0,1,1), alpha=0.05); print(est)
#str(monarch.fit)
#monarch.fit$logliks
plot(monarch.fit, which=1)
plot(monarch.fit, which=3)

#Try a different estimator
# monarch.fit1 <- chngptm(formula.1= y~1, formula.2=~year, monarchdata.df,
#                         type="segmented", family="gaussian", est.method="smoothapprox", var.type="bootstrap", save.boot=TRUE)
# summary(monarch.fit1)
# est1 <- lincomb(monarch.fit1, comb=c(0,1,1), alpha=0.05); print(est1)
# 
#Note: warning that fastgrid2 is a better for linear regression
#
# plot(monarch.fit1, which=1)
# plot(monarch.fit1, which=3)

#Yet a different estimator
monarch.fit2 <- chngptm(formula.1= y~1, formula.2=~year, monarchdata.df,
                        type="segmented", family="gaussian", est.method="fastgrid2", var.type="bootstrap", save.boot=TRUE)
summary(monarch.fit2)
est2 <- lincomb(monarch.fit2, comb=c(0,1,1), alpha=0.05); print(est2)
chngpt.test(
  formula.null = y ~ 1,
  formula.chngpt =  ~ year,
  data = monarchdata.df,
  type = "segmented",
  family = "gaussian",
  test.statistic = "lr",
  p.val.method = "param.boot"
)

#str(monarch.fit2)  #AIC = 45.3


###Comparing the segmented model to the step model
monarch.fit2$best.fit$aic

deltaAIC <- monarch.step$best.fit$aic - monarch.fit2$best.fit$aic
pAIC <- exp(-deltaAIC/2); print(pAIC)

(1-pAIC)/pAIC

####Plotting

plot(monarch.fit2, which=1)
plot(monarch.fit2, which=3)
plot(monarch.fit2, which=2)
str(monarch.fit2)
monarch.fit2$logliks

plot(monarch.fit2, xlim=c(1994, 2019), pch=19, lwd=5, col="red", which=1)
abline(v=2019, lty=2, col="black")
abline(h=log(6), lty=3, col="gray")

plot(monarch.fit2, xlim=c(1994, 2020), pch=19, lwd=5, col="red", which=1)
abline(v=2019, lty=2, col="black")
abline(v=2020, lty=2, col="black")
abline(h=log(6), lty=3, col="gray")

plot(monarch.fit2, xlim=c(1994, 2021), pch=19, lwd=5, col="red", which=1)
abline(v=2019, lty=2, col="black")
abline(v=2020, lty=2, col="black")
abline(v=2021, lty=2, col="black")
abline(h=log(6), lty=3, col="gray")

par(mfrow=c(1,1))

#Test whether changepoint exists
test <- chngpt.test(formula.null= y~1, formula.chngpt=~year, monarchdata.df,
                 type="segmented", family="gaussian")
print(test)
plot(test)


#plotting
library(ggplot2)
library(ggpubr)
library(ggExtra)

monarchdata.df$group <- ifelse(monarchdata.df$year < 2014, "Pre-2014", "Post-2014")  #Based on changepoint identified above

# p <- ggscatter(monarchdata.df, x = "year", y = "y",
#           add = "reg.line",                         # Add regression line
#           conf.int = TRUE,                          # Add confidence interval
#           color = "group", palette = c("darkorange3", "steelblue4"),         # Color by groups 
#           size = 5, alpha = 0.8,
#           shape = 19,
#           xlim=c(1994,2018), label = "year", repel = TRUE
# )  + theme(legend.position="bottom")
# 
# p

parOrig <- par()

# par(bg=NA)
p <- ggplot(monarchdata.df)  +
  geom_smooth(method = lm,
              aes(
                x = year,
                y = y,
                fill = group,
                color = group
              ),
              colour = "black") +
  geom_point(
    aes(x = year, y = y),
    shape = 19,
    size = 5,
    alpha = 0.8,
    color = "black"
  ) +
  #  geom_histogram(data=bootyears, aes(x=year), bins=25, colour="light gray") +
  theme_classic(base_size = 15) +
  scale_fill_manual(values = c('#999999', '#E69F00')) +
  theme(legend.position = "none") +
  scale_x_continuous(
    name = "Year",
    limits = c(1994, 2018),
    breaks = pretty(monarchdata.df$year, n = 10)
  ) +
  scale_y_continuous(
    name = expression(paste(log[e](Area (
      ha
    )))),
    limits = c(min(y - 0.3), max(y + 0.3)),
    breaks = pretty(monarchdata.df$y, n = 10),
    sec.axis = sec_axis( ~ 150 * ., labels = NULL, name = "Frequency
                         ")
    )

p


bootyears <- as.data.frame(monarch.fit2$vcov$boot.samples[,4])
colnames(bootyears) <- "year"
# p.boot <- ggplot(bootyears, aes(year))  + 
#   geom_histogram() +
#   theme_classic(base_size = 15) + 
#   theme(legend.position = "none", axis.text.y = element_blank(), axis.text.x = element_blank()) +
#   scale_x_continuous(name="", limits=c(1994, 2018)) +
#   scale_y_continuous(name="")
# 
# p.boot


#############
#How many more years of positive increase before increase can be said to be significant?

#Scenario 1, what if the mean increase from the post-2014 period continued 1 additional year?
monarchdata.pred1 <-
  rbind(monarchdata.df, c(2019, monarchdata.df[21, 2] + log(exp(est2) * 5)))


monarch.pred1 <-
  chngptm(
    formula.1 = y ~ 1,
    formula.2 =  ~ year,
    monarchdata.pred1,
    type = "segmented",
    family = "gaussian",
    est.method = "fastgrid2",
    var.type = "bootstrap",
    save.boot = TRUE
  )
summary(monarch.pred1)
pred1 <- lincomb(monarch.pred1, comb=c(0,1,1), alpha=0.05); print(pred1)  #print(est2)

plot(monarch.pred1, which=1)
plot(monarch.pred1, which=3)

#Conclusion: 1 additional year of growth at the post-2014 mean rate of change will be sufficient for declaring a significant positive trend


#Scenario 2, how low can the 2019-2020 population be and still return a post-2014 rate of change that is positive and significant?
monarchdata.pred2 <- rbind(monarchdata.df, c(2019, monarchdata.df[21,2]+0.667*log(exp(est2)*5))) 


monarch.pred2 <- chngptm(formula.1= y~1, formula.2=~year, monarchdata.pred2,
                         type="segmented", family="gaussian", est.method="fastgrid2", var.type="bootstrap", save.boot=TRUE)
summary(monarch.pred2)
pred2 <- lincomb(monarch.pred2, comb=c(0,1,1), alpha=0.05); print(pred2)  #print(est2)

exp(monarchdata.df[21,2]+0.667*log(exp(est2)*5)) #4.00

#Conclusion, if next year's observed estimate is >=4.0, then this will mean the period from 2014-2019 continues to exhibit positive growth
#Any less than 4.0 and the post-2014 period cannot credibly be described as growing.

#####Scenario, if 2019 = 4.0 but 2020 = 2.5, what is the conclusion?
#monarchdata.pred1 <- rbind(monarchdata.df, c(2019, monarchdata.df[21,2]+log(exp(est2)*5))
monarchdata.pred4 <- rbind(monarchdata.df, c(2019, monarchdata.df[21,2]+log(exp(est2)*5)), c(2020, log(2.5)))

monarch.pred4 <- chngptm(formula.1= y~1, formula.2=~year, monarchdata.pred4,
                         type="segmented", family="gaussian", est.method="fastgrid2", var.type="bootstrap", save.boot=TRUE)
summary(monarch.pred4)
pred4 <- lincomb(monarch.pred4, comb=c(0,1,1), alpha=0.05); print(pred4)  #print(est2)
plot(monarch.pred4, which=1)
plot(monarch.pred4, which=3)


#monarchdata.pred2 <- rbind(monarchdata.pred1, c(2020, monarchdata.df[26,2]+est2)) #Unnecessary given that 1 additional year makes the trend post-2014 significant

#Scenario 2, what if the increase was only half as much?
monarchdata.pred2 <- rbind(monarchdata.df, c(2019, monarchdata.df[25,2]+0.5*est2)) 


monarch.pred2 <-
  chngptm(
    formula.1 = y ~ 1,
    formula.2 =  ~ year,
    monarchdata.pred2,
    type = "segmented",
    family = "gaussian",
    est.method = "fastgrid2",
    var.type = "bootstrap",
    save.boot=TRUE)
summary(monarch.pred2)
pred2 <- lincomb(monarch.pred2, comb=c(0,1,1), alpha=0.05); print(pred2)

plot(monarch.pred2, which=1)
plot(monarch.pred2, which=3)

#Scenario 3
monarchdata.pred3 <- rbind(monarchdata.df, c(2019, monarchdata.df[25,2]+0.75*est2)) 


monarch.pred3 <-
  chngptm(
    formula.1 = y ~ 1,
    formula.2 =  ~ year,
    monarchdata.pred3,
    type = "segmented",
    family = "gaussian",
    est.method = "fastgrid2",
    var.type = "bootstrap",
    save.boot=TRUE)
summary(monarch.pred3)
pred3 <- lincomb(monarch.pred3, comb=c(0,1,1), alpha=0.05); print(pred3)

plot(monarch.pred3, which=1)
plot(monarch.pred3, which=3)


#Scenario 4
monarchdata.pred4 <- rbind(monarchdata.df, c(2019, monarchdata.df[25,2]+0.95*est2)) 


monarch.pred4 <-
  chngptm(
    formula.1 = y ~ 1,
    formula.2 =  ~ year,
    monarchdata.pred4,
    type = "segmented",
    family = "gaussian",
    est.method = "fastgrid2",
    var.type = "bootstrap",
    save.boot=TRUE)
summary(monarch.pred4)
pred4 <- lincomb(monarch.pred4, comb=c(0,1,1), alpha=0.05); print(pred4)

plot(monarch.pred4, which=1)
plot(monarch.pred4, which=3)


# 
# ###sparklines used in manuscript
# library(sparkline)
# yearspark <- seq(from=1994, to=2018, by=1)
# stepspark <- c(6,6,6,6,6,6,6,6,6,6,6,6,6,6,6,6,2,2,2,2,2,2,2,2,2)
# segspark <- c(3.21, 3.10, 2.95, 2.81, 2.67, 2.54, 2.40, 2.27, 2.13, 2.00, 1.87, 1.73, 1.60, 1.47, 1.33, 1.20, 1.06, 0.93, 0.80, 0.67, 1.04, 1.41, 1.78, 2.15, 2.52)
# sparkmonarch <- as.data.frame(cbind(yearspark, stepspark,segspark))
# 
# sparkline(sparkmonarch$stepspark, line.pars = gpar(lwd=5), buffer = unit(10, "lines"))
# sparkline(sparkmonarch$segspark, line.pars = gpar(lwd=5), buffer = unit(10, "lines"))
# 

  

########## Figure 1
y<- log(c(7.81,12.61,18.19,5.77,5.56,9.05,2.83,9.36,7.54,11.12,2.19,5.91,
          6.87,4.61,5.06,1.92,4.02,2.89,1.19,0.67,1.13,4.01,2.91,2.48,6.05))
year <-seq(1994,2018)

plotdata <- cbind.data.frame(year, y)

library(ggplot2)


gg9414 <- ggplot(plotdata, aes(year,y))  +  geom_point() +
  geom_smooth(data=plotdata[plotdata$year>=1994 & plotdata$year<=2014,],method = "lm") 


gg_data9414 <- ggplot_build(gg9414)
#str(gg_data9414)
head(gg_data9414$data[[2]])
gg9414.2 <- gg_data9414$data[[2]]


gg9418 <- ggplot(plotdata, aes(year,y))  +  geom_point() +
  geom_smooth(data=plotdata[plotdata$year>=1994 & plotdata$year<=2018,],method = "lm") 


gg_data9418 <- ggplot_build(gg9418)
#str(gg_data9418)
head(gg_data9418$data[[2]])
gg9418.2 <- gg_data9418$data[[2]]

gg1318 <- ggplot(plotdata, aes(year,y))  +  geom_point() +
  geom_smooth(data=plotdata[plotdata$year>=2013 & plotdata$year<=2018,],method = "lm") 


gg_data1318 <- ggplot_build(gg1318)
#str(gg_data1318)
head(gg_data1318$data[[2]])
gg1318.2 <- gg_data1318$data[[2]]


plotgg <- ggplot(plotdata, aes(year,y))  +
  theme_classic(base_size = 15) + 
  geom_smooth(data=plotdata[plotdata$year>=1994 & plotdata$year<=2014,],method = lm, se=FALSE, linetype="ff",
              fill='#999999', color='black') + 
  geom_smooth(data=plotdata[plotdata$year>=1994 & plotdata$year<=2018,],method = lm, se=T, 
              fill='lightblue1', color='black') + 
  geom_smooth(data=plotdata[plotdata$year>=2013 & plotdata$year<=2018,],method = lm, se=T, 
              fill='#E69F00', color='black') +
  geom_line(data = gg9414.2, aes(x = x, y = ymin), size = 0.02, linetype="dashed") +
  geom_line(data = gg9414.2, aes(x = x, y = ymax), size = 0.02, linetype="dashed") +
  geom_line(data = gg9418.2, aes(x = x, y = ymin), size = 0.02, linetype="solid") +
  geom_line(data = gg9418.2, aes(x = x, y = ymax), size = 0.02, linetype="solid") +
  geom_line(data = gg1318.2, aes(x = x, y = ymin), size = 0.02, linetype="solid") +
  geom_line(data = gg1318.2, aes(x = x, y = ymax), size = 0.02, linetype="solid") + 
  annotate("text", x = c(2015.5, 2012, 2007), y = c(2.5, 1.6, 0.4), 
           label = c("Post change","Full series", "Semmens et al. series")) +
  geom_point(shape = 19, size = 5, color = "black") + 
  scale_x_continuous(
    name = "Year",
    limits = c(1994, 2018),
    breaks = pretty(year, n = 10)
  ) +
  scale_y_continuous(
    name = expression(paste(log[e](Area (
      ha
    )))),
    limits = c(min(y - 0.5), max(y + 0.5)),
    breaks = pretty(y, n = 10),
  ) 

plotgg

############
#Extinction Risk
############


###     Original Semmens et al. data (to 2014)

#See accompanying file 


###     Semmens et al. model for full timeseries

#see accompanying file monarchSS extinction_cloning_obsSmaller3add2018.R

                         

###     Count-Based PVA for post-change point period


#########################################

# ALTERNATIVE ANALYSIS using data from the post-changepoint years to represent current conditions 
year = seq(2014,2018)
N= c(1.13,4.01,2.91,2.48,6.05)

#########################################

#check that the data are entered correctly
a= length(year)
a
year
b=length(N)
b
N

# calculate annual growth rates
# recall that the intrinsic rate of increase (r)= ln(lambda), lambda = e^r
# and of course we can't get lambda for the lastyear
ln.lambda=log(N[-1]/N[-length(N)]) 

# just renaming the growth rate variable to help keep things clear
r=ln.lambda


##############Summary of the data and inspection
r
summary(r)
boxplot(r)
hist(r)
geomean=mean(r)
#examine the geometric mean intrinsic rate of increase
geomean

# and the variance and standard deviation are calculated from the ln(lambda) values, 
# because the growth rates (lambdas) are lognormally distributed while the r values are
# normally distributed
var.r=var(r)
sd.r=sd(r)
var.r
sd.r


# examine the actual population dynamics that were observed
plot(N~year, xlab = "Year", ylab = "Population size", pch = 19, ylim=c(0,20),
     type="b", main="Monarch Butterfly population size over years",cex.lab=1.5,
     cex=1,cex.axis=1.5)
############## 


# add a trajectory for simple exponental growth over the same period, with
# the observed geomean for r.  
lines(year,N[1]*exp(geomean)^(0:(length(N)-1)),pch=19,col='red',type="b",cex=0.8)
legend(2015,18,c("Empirical data","Exponential Model prediction"),lty=c(1,1),col=c("black","red"),pch=c(19,19), bty="y")
############

################

N0 = N[5]   # setting initial population size for the projections equal to original pop size in the data


# Define a threshold population size that we consider unacceptable.  For true local extirpation, this would be zero. 

threshold=0.25             

# Set the number of years to project forward (checking match to 5 years of data, plus 5 years projection)

#project= n, need to match this in column extracted to plot in hist() below
project = 6

# Set the of model iterations to perform

runs=1000  

# Create an empty matrix to hold the output (population size with stochastic exponential growth), for the specified number of years (as rows) and iterations (as columns).  

stoch.pop=matrix(NA,project,runs)       
stoch.pop[1,]=N0


# two nested loops to create stochastic population sizes

for (i in 1:runs){					 # looping over 1000 runs of the stochastic model
  for (t in 2:project){			# and looping over n years of projection within each of 1000 runs
    lambda=exp(rnorm(1,geomean,sd.r))		# draw a value of lambda from a lognormal distribution
    stoch.pop[t,i]=stoch.pop[(t-1),i]*lambda	# and project one time step from the current pop size
    if(stoch.pop[t,i]<=threshold) break  		# leave the loop if pop <= threshold
  }
}

# the stochastic output 

stoch.pop[1:6, 1:10]
stoch.pop[1:6, 991:1000]

# the frequency distribution for population sizes in 2023, for all 1000 iterations

hist(stoch.pop[6,], xlab = 'Population Size', font = 2, font.lab = 2, main = 'Monarch butterfly population, 2023')

############### PERCENTAGE OF RUNS UNDER RISK THRESHOLD POPULATION SIZE 

percentage.under<-(runs-length(which(stoch.pop[project,]>=0.25)))/runs*100 
percentage.under  # the number of runs with population size lower than threshold
#################

############### MEAN YEAR WHEN POPULATION SIZE WENT BELOW THRESHOLD, FOR THOSE RUNS WITH A PSEUDO-EXTINCTION
time<-NULL				## create an empty vector to hold results
for (i in 1:runs){			          ## loop through all the iterations of the simulation
  t<-max(which(stoch.pop[,i]>0)) 	## Find the maximum time when Population was >0 (as opposed to remaining NA) for each run
  time<-c(time,t)			## Add this value to the end of a vector storing the times
}					## end the loop
time.under<-time[which(time<10)]
hist(time.under,nclass=20, xlab = 'Year at which pseudoextinction occurred')
abline(v=median(time.under),lw=3)
#################


##################################### Environmental stochasticity

project=6 
runs=1000  
stoch.pop=matrix(NA,project,runs)

## Divide r empirical data into "bad years (r<0)" and "good years (r>0)"

geomean.bad<-mean(r[which(r<0)])
geomean.good<-mean(r[which(r>0)])

stoch.pop[1,]=N0

# loop to create stochastic population sizes
for (i in 1:runs){					             # looping over 1000 runs of the stochastic model
  for (t in 2:project){					          # and looping for n years of projection    
    geomean=sample(x=c(geomean.bad,geomean.good),size=1,prob=c(0.5,0.5),replace=T)  
    #Sample 1 number from a vector x with probabilities prob, here 50/50 for good/bad
    lambda=exp(rnorm(1,geomean,sd.r))		        
    # draw a value of lambda from a lognormal distribution with the specified mean
    stoch.pop[t,i]=stoch.pop[(t-1),i]*lambda	  
    # and project one time step from the current pop size
    if(stoch.pop[t,i]<=threshold) break  			  # leave the loop if pop <= threshold
  }
}

# examine the stochastic output (note that this is huge, 6 x 1000) so we only look at a bit of it
stoch.pop[1:6,1:10]
stoch.pop[1:6, 991:1000]

percentage.under<-(runs-length(which(stoch.pop[project,]>=0.25)))/runs*100 
percentage.under
########################################################


# Plot the projected results

stoch.pop.mean=apply(stoch.pop,1,mean, na.rm=T)
log.pop.sd  =apply(log(stoch.pop),1,sd, na.rm=T)
ucl =exp(log(stoch.pop.mean)+1.96*log.pop.sd)     #upper confidence limit
lcl =exp(log(stoch.pop.mean)-1.96*log.pop.sd)     #lower confidence limit
plot(1:project,stoch.pop.mean,'b',pch = 19, col = 'blue', ylim=c(0,18.19),xlab='Years from 2017', ylab='Population Size')
lines(1:project,lcl,'l', col = 'blue')
lines(1:project,ucl,'l', col  = 'blue')


#################################################
# same again with log Y axis so that the upper CL does not obscure other patterns
plot(1:project,stoch.pop.mean, log = "y", type = 'b', pch = 19, cex=2, col = 'black', ylim=c(1,max(ucl)), xlab='Years from 2017',
     ylab='log(Area [ha])')
lines(1:project,lcl,'l',  col = 'black')
lines(1:project,ucl,'l',  col  = 'black')
#points(1:length(year), N, pch = 1, col = 'red', bg = 'red', cex=2)
legend(1,10000, c("Projection with 95% CL", "Data"), lty = 1, col = c('black', 'red'), pch =c(19,1),cex=1,bt="n")
abline(h=18.19, lty=2, col="black") #with 18.19ha identified as upper limit of population


# Simple population-projection population viability analysis using popbio package

library(popbio)  

set.seed(4)
# number of projections, i.e. the number of simulated propoualtion growth trajectories
reps = 1000

#set of growth rates to sample with replacement for growth at each time step
lambda=(N[-1]/N[-length(N)])
K = 18.19  #has a potentially large effect on probability of extinction, but is used to control for the ridiculous growth that occurs for an uncapped population
damp = 1 - (N[-1]/K)
dd.lambda = lambda*damp
lambda
damp
dd.lambda 

final.N<-stoch.projection(as.list(dd.lambda), N[5], nreps=reps, tmax = 5) #tmax = 2023; alternatively, nmax=18.19
final.N

###############################
ext = 0
ext.threshold = 0.25  #  zero the counter for number of extinctions before tallying extinctions, quasi-extinction threshold set to 0.25 ha

#loop to tally up the number of cases where N.final is < ext.threshold
for (i in 1:length(final.N))
{if (final.N[i] < ext.threshold) ext = ext+1
}
ext  # number of cases where N.final is < ext.threshold
prob.ext = ext/reps
prob.ext  # proportion of cases where N.final is < ext.threshold, i.e. Prob{pseudoextinction}


####
final.N<-stoch.projection(as.list(lambda), N[5], nreps=reps, tmax = 5) #tmax = 2023; alternatively, nmax=18.19
final.N
summary(final.N)

###############################
ext = 0
ext.threshold = 0.25  #  zero the counter for number of extinctions before tallying extinctions, quasi-extinction threshold set to 0.25 ha

#loop to tally up the number of cases where N.final is < ext.threshold
for (i in 1:length(final.N))
{if (final.N[i] < ext.threshold) ext = ext+1
}
ext  # number of cases where N.final is < ext.threshold
prob.ext = ext/reps
prob.ext  # proportion of cases where N.final is < ext.threshold, i.e. Prob{pseudoextinction}


###alternative diffusion approximation
countCDFxt(mu=geomean, sig2=var.r, nt=5, Nc=6.06, Ne=0.25)
options(scipen = 999)
#options(scipen = 0)
#in year 5 (2023)  Gbest = 0.00001134498332 Glo = 0.00000000000000000008788863542 Gup = 0.14773653


########
matrix(rnorm(500 * 20, mean = 6, sd = sqrt(1.52)), nrow = 500)


e <- list()
f <- list()
g <- list()
h <- list()
j <- list()

for (i in 1:500) {
  e[i] <- list(rnorm(n = 3, mean = 7, sd = sqrt(1.52)))
  f[i] <- list(rnorm(n = 3, mean = 6.85, sd = sqrt(1.52)))
  g[i] <- list(rnorm(n = 5, mean = 7, sd = sqrt(1.52)))
  h[i] <- list(rnorm(n = 6, mean = 7, sd = sqrt(1.52)))
  j[i] <- list(rnorm(n = 7, mean = 7, sd = sqrt(1.52)))
}

esap <- sapply(e,mean)
mean(esap)
eerror <- qnorm(0.975)*sd(esap)/sqrt(3)
eleft <- mean(esap)-eerror
eright <- mean(esap)+eerror
eleft
eright

fsap <- sapply(f,mean)
mean(fsap)
ferror <- qnorm(0.975)*sd(fsap)/sqrt(3)
fleft <- mean(fsap)-ferror
fright <- mean(fsap)+ferror
fleft
fright
